# Patient-derived xenograft studies of fumarate hydratase (FH)-deficient uterine leiomyoma subtype

**DOI:** 10.1101/2022.07.27.501688

**Authors:** Takuya Kajimura, Vanida Ann Serna, Emma Amurgis, Michael Blumenfeld, Takeshi Kurita

## Abstract

Uterine leiomyoma (LM) is the most common benign gynecological tumor in premenopausal women. Our previous patient-derived xenograft (PDX) studies established that 17ß-estradiol (E2) and progesterone (P4) stimulate the growth of the two most prevalent subtypes, MED12 mutant (MED12-LM) and HMGA2 overexpressing LMs (HMGA2-LM), via proliferation and hypertrophy of smooth muscle tumor cells (SMTCs). In addition, tumor-associated fibroblasts (TAFs) that do not carry *MED12* mutations also contribute to the growth of MED12-LM by secreting extracellular matrix (ECM) proteins. In this study, we investigated the growth control of the fumarate hydratase (FH) deficient LM (FH-LM) subtype, utilizing the PDX model. We identified an FH-negative case with conventional leiomyoma histology. The overexpression of aldo-keto reductase family 1 member B10 (AKR1B10) confirmed the FH deficiency. Like MED12-LM, FH-LM comprised two major cell types: 54.4% SMTCs and 43.3% TAFs. Furthermore, the TAFs expressed FH. The FH-LM PDXs grew in response to E2 and P4 via proliferation and hypertrophy of SMTCs, similar to MED12-LM and HMGA2-LM. While E2 alone did not stimulate growth, E2 was essential for sensitizing FH-deficient SMTCs to P4 by upregulating progesterone receptor (PGR). Our current study established that the growth of the three most prevalent LM subtypes, MED12-LM, HMGA2-LM, and FH-LM, depends on E2 and P4. Thus, selective progesterone receptor modulators (SPRMs) should be an effective treatment option for most symptomatic LM patients.

## Introduction

Uterine leiomyoma (LM), also known as uterine fibroid, is a benign smooth muscle tumor of the myometrium (MM) with a cumulative incidence of approximately 70% [1, 2]. Recent transcriptome and genomic analyses identified four major LM subtypes with distinct molecular profiles, *MED12* mutant (MED12-LM), HMGA2 overexpressing (HMGA2-LM), fumarate hydratase (*FH*) deficient (FH-LM), and *COL4A5*-*COL4A6*-deletion subtypes (COL4A5/6-LM) [3, 4]. Additionally, there are LMs that do not bear any of the major 4 subtype genetic alterations (quadruple-negative-LMs). MED12-LM and HMGA2-LM are the two most common subtypes accounting for approximately 70% and 15% of all LM cases, respectively. Our previous patient-derived xenograft (PDX) studies established that 17ß-estradiol (E2) and progesterone (P4) stimulate the growth of these two major LM subtypes via proliferation and hypertrophy of smooth muscle tumor cells (SMTCs) [5-7]. Meanwhile, the cellular composition of MED12-LM and HMGA2-LM is distinctive: tumor-associated fibroblasts (TAFs), which do not carry driver mutations, account for >40% of the cell population in MED12-LM but <10% in HMGA2-LM. In MED12-LM, TAFs also contribute to tumor growth by secreting extracellular matrix (ECM) [7, 8]. While the unique genetic mutations and gene expression profiles suggest distinctive biology, the growth regulation of LM subtypes other than MED12-LM and HMGA2-LM are currently unknown.

FH-LM is the third most prevalent LM subtype occurring in 0.4 - 2.6% of all LM cases [9-12]. FH is a tricarboxylic acid (TCA) cycle enzyme that catalyzes the reversible stereospecific hydration of fumarate to L-malate, and its loss of function is associated with tumorigenesis [13]. However, the exact mechanisms of how FH-deficiency drives LM formation remain unknown. FH-LM is often associated with Hereditary Leiomyomatosis and Renal Cell Cancer (HLRCC), a hereditary syndrome predisposed to cutaneous leiomyoma, LM, and renal cell cancer due to germline mutations of *FH*. However, recent studies suggest that sporadic FH-LM cases in non-HLRCC carriers are as common as HLRCC-associated FH-LM cases [12, 14]. In addition, neither mutation analysis nor FH immunohistochemistry is sufficient to identify FH-LM because the loss of FH protein can occur independently of *FH* mutations, and a small proportion of FH-LMs retain FH protein [12, 14, 15]. Thus, the true prevalence of FH-LM may be higher than current estimates. Accordingly, understanding the pathogenesis of FH-LM subtype is crucial. Primary cell culture has been a standard research model of LMs. However, SMTCs cannot be maintained in 2D cell culture [16, 17]. In addition, *in vitro* models, including 3D spheroid/organoid cultures, do not replicate the hormone-dependent growth of LM cells [18]. Thus, in this manuscript, we investigated the growth control of FH-LM, utilizing a patient-derived xenograft (PDX) model, which faithfully replicates the hormone-dependent growth of LM [5, 19].

## Materials and Methods

### Collection and characterization of LM cases

The acquisition and research use of surgical specimens were approved by the Institutional Review Boards of the Ohio State University and Northwestern University. LM and MM samples were obtained from hysterectomy or myomectomy patients with prior written informed consent and delivered to research personnel within 5 hours of surgical removal. The MED12-LM and HMGA2-LM subtypes were identified as previously described [7]. To determine *MED12* genotype, genomic DNA was extracted from LM and MM samples, and the sequence of *MED12* exon 1, intron 1, and exon 2 was determined by the Sanger sequencing of PCR products. To amplify *MED12* from exon 1 to exon 2, we used two primer sets, 5’-gtcggtattgtccgatggtt-3’ (forward) and 5’-gtcagtgcctcctcctagg-3’ (reverse) and 5’-ggtggctgggaatcctagtg-3’ (forward) and 5’-ccctataagtcttcccaaccca-3’ (reverse). LM cases were classified as HMGA2-LMs when >50% of cells showed intense nuclear staining in HMGA2 IHC. In this study, the genomic DNA was extracted from the formalin-fixed paraffin-embedded tissues. However, sequence analysis of *FH* and *COL4A5*-*COL4A6* loci was impossible due to the low yield and quality of genomic DNA. Accordingly, FH-LM and COL4A5/6-LM were identified by the loss of FH and COL4A5 proteins utilizing IHC.

### PDX experiment

The procedures for PDX model preparation have been described previously [19]. Twenty-five PDXs were prepared from a single FH-LM sample and grafted into 12 adult NSG (NOD.Cg-*Prkdc*^*scid*^ *Il2rg*^*tm1Wjl*^/SzJ) female mice (Jackson Laboratory, Bar Harbor, ME, USA). All host mice were ovariectomized and subcutaneously implanted with a 70 mg slow-releasing pellet containing E2 and P4 (E2P4) [19]. Four weeks later, one group of mice was euthanized to collect PDXs, and E2P4 pellets were removed from the remaining hosts. These host mice were divided into 3 groups (N = 2) and implanted with no hormone (NH), a 70 mg slow-releasing E2 pellet, or a new E2P4 pellet (E2P4). They were euthanized 2 weeks after hormone pellet replacement to collect PDXs. After measuring the tumor volume [19], PDXs were fixed with Modified Davidson’s fixative solution (Electron Microscopy Sciences, Hatfield, PA) overnight and processed into paraffin blocks processed for histologically analysis, as previously described [20]. This experiment was carried out in strict accordance with the recommendations in the Guide for the Care and Use of Laboratory Animals of the National Institutes of Health. The protocol was approved by the Institutional Animal Care and Use Committee (IACUC) of the Ohio State University (Protocol Number: 2014A00000060). All surgery was performed under ketamine and xylazine anesthesia, and all efforts were made to minimize suffering.

### Immunostaining

IHC with DAB (3,30-diaminobenzidine) and immunofluorescence (IF) were performed following the methods previously described [21] with minor modifications. Paraffin blocks were sectioned at 5 µm and mounted on ASI Supreme Frosted Glass Microscope Slides (Alkali Scientific, Fort Lauderdale, FL). Slides were pre-heated on a slide warmer at 60 °C for >15 min and deparaffinized through the series of xylene and ethanol. For hematoxylin and eosin (H&E) staining, slides were stained with SelecTech Hematoxylin/Eosin Staining System (Leica Biosystems, Buffalo Grove, IL). For immunostaining, slides were emersed in 10mM sodium citrate buffer (pH 6.0) containing 0.05% Tween 20 and heated for 30 min in an Electric Pressure Cooker. Tissue sections were separated by drawing a circle around with a PAP pen (Daido Sangyo, Tokyo, Japan) and incubated with a blocking buffer (2% donkey serum, 1% BSA, 0.1% Cod fish gelatin, 0.1% TritonX100, 0.05% sodium azide, 0.05 % Tween 20, 10mM PBS) at a room temperature (RT, ∼20 – 23 °C) for 60 min, followed by incubation with a primary antibody at 4 °C overnight. The following primary antibodies were used at indicated dilutions: anti-FH (1:200,10966-1-AP, Proteintech, Rosement, IL), anti-HMGA2 (1:800, #8179) and anti-vimentin (VIM) (1:200, #9856) (Cell Signaling Technologies, Danvers, MA), anti-MKI67 (1:100, ab92742), anti-calponin (1:100, ab197639), anti-TOMM20 (1:200, ab56783) and anti-ACTA2 (1:500, ab781) (Abcam, Boston, MA), anti-COL4A5 (1:1000, PA5-119042), anti-ARK1B10 (1:1000, PA5-22036) (Thermo Scientific, Waltham, MA), anti-ESR1 (1:100 RM9101-S, Lab Vision), anti-MED12 (1:50, HPA003184, Sigma-Aldrich, St. Louis, MO) and anti-PGR (1:200, A0098, Agilent Technologies, Santa Clara, CA). For IHC, biotinylated anti-rabbit IgG (H+L) (1:800, 711-066-152, Jackson ImmunoResearch, West Grove, PA) was used as the secondary antibody in combination with streptavidin-horseradish peroxidase (1:400, 016-030-084, Jackson ImmunoResearch). IHC stained sections were counter-stained with Hematoxylin 560 MX (Leica Biosystems). For IF, the primary antibody was detected utilizing Alexa-Fluor594 anti-mouse IgG (H+L) (1:1000, 715-586-151, Jackson ImmunoResearch) and Alexa-Fluor488 anti-rabbit IgG (H+L) (1:100, 711-546-152, Jackson ImmunoResearch). For IF, the nucleus was stained with Hoechest 33258 (1:10000, Sigma-Aldrich). Micrographs were captured using a BZ-9000 microscope (Keyence, Itasca, IL).

### Morphometric analysis

The morphometric analyses were performed as previously described [7]. For the analysis of original tumors, at least 3 pieces sampled from different parts of the tumor were included in the analysis.

The cellular composition of smooth muscle cells (SMCs) versus non-SMCs was determined by counting the nuclei of calponin (SMC marker)-positive and negative cells (total >200 cells per section x >3 sections per sample) in tissue sections stained for calponin and vimentin (VIM). The concentration of fibroblasts was determined by counting non-vascular cells positive for VIM but negative for calponin. In the cell composition analysis of the FH-LM case, 5 pieces dissected from different parts of the original tumor were considered independent samples.

The MKI67 labeling indices were determined by manually counting positive and negative cells for ACTA2 (SM marker) and MKI67 in double IF-stained tissue sections. At least 300 cells per sample and 1,000 cells per group were counted blindly. SMTC size was calculated as total ACTA2-positive pixel number divided by the number of ACTA2-positive cells (ACTA2-positive area per SMTC cell). In this analysis, at least 300 cells per sample and 1,000 total cells per group were counted blindly. Cell density was determined as the number of nuclei within target areas (>5 mm^2^ per group and >1 mm^2^ per sample). For each sample, at least 500 total cells from five fields (40x magnification) were counted blindly. We used analysis of variance (ANOVA) to compare more than two groups, and p<0.05 was considered significant. Data were presented as mean values with standard deviation (SD).

## Results

### Analysis of archived LM samples used to generate PDXs

We determined the subtypes of 17 archived LM cases used for PDX studies, including 12 cases analyzed in our previous studies [5-8, 19] (Table 1). All 17 cases showed conventional LM histology (Fig 1A). Among these samples, 9 cases (52.9%) harboring *MED12* mutations were classified as MED12-LM (Table 1). Nevertheless, there was no evident difference in the expression pattern of MED12 among all LM cases and MM, as assessed by IHC (Fig 1B). Among 17 cases, 4 LMs (23.5%) were classified into HMGA2-LM, presenting an elevated level of HMGA expression (Fig 1B). These 4 HMGA2-LM cases were used in our previous PDX studies [7]. Additionally, IHC screening identified a LM case negative for FH (Fig 1B). Three cases (17.6%) were negative for *MED12* mutations and HMGA2 overexpression but positive for FH expression (Table1). All 17 LM cases, including the 3 LMs of unknown subtype, were positive for COL4A5 by IHC (Fig 1B).

**Table 1.** Histological character of LM tumors

| case. ID | LM subtype | MED12 mutant | HMGA2 overexpression | FH expression |
| --- | --- | --- | --- | --- |
| 1 | HMGA2-LM | WT | + | + |
| 2 | MED12-LM | c.130G>C | - | + |
| 3 | MED12-LM | c.116T>C | - | + |
| 4 | MED12-LM | c.131G>T | - | + |
| 5 | HMGA2-LM | WT | + | + |
| 6 | MED12-LM | c.130G>C | - | + |
| 7 | HMGA2-LM | WT | + | + |
| 8 | MED12-LM | c.131G>T | - | + |
| 9 | MED12-LM | c.131G>A | - | + |
| 10 | MED12-LM | c.131G>A | - | + |
| 11 | HMGA2-LM | WT | + | + |
| 12 | unknown | WT | - | + |
| 13 | FH-LM | WT | - | - |
| 14 | unknown | WT | - | + |
| 15 | unknown | WT | - | + |
| 16 | MED12-LM | c.131G>C | - | + |
| 17 | MED12-LM | c.131G>T | - | + |

**Figure. 1.**
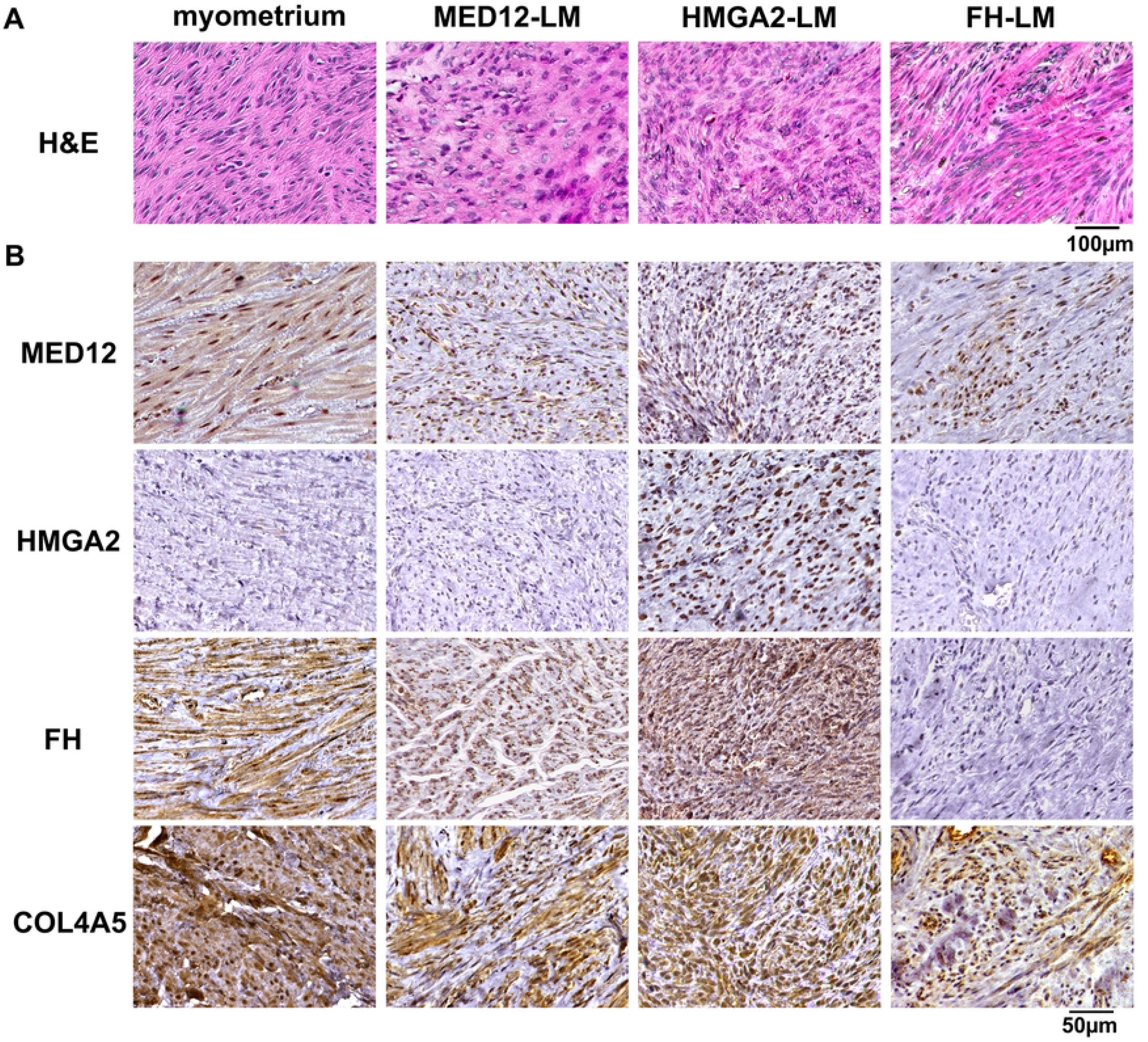
Histological analysis of MM and LM samples. Representative images of H&E (**A**) and IHC (**B**) stained MM and LM subtypes.

### Histological characteristics of FH-LM

The FH-LM case showed the morphological characteristics of FH-deficient LMs [11, 22], including staghorn vessels (Fig 2Aa, Arrow), and enlarged nuclei with pseudoinclusions (Fig 2Ac) [23]. IF assay for calponin (SMC marker) and VIM showed that the FH-LM contained a substantial concentration (43.3%) of TAFs (Fig 2B). The SMTC concentrations were not statistically different among FH-LM (54.4%), MED12-LM (54.3%), and MM (61.2%), whereas HMGA2-LM contained a significantly higher concentration of SMTCs (91.4%) than other LM subtypes and MM (Fig 2C) (P<0.001).

**Figure. 2.**
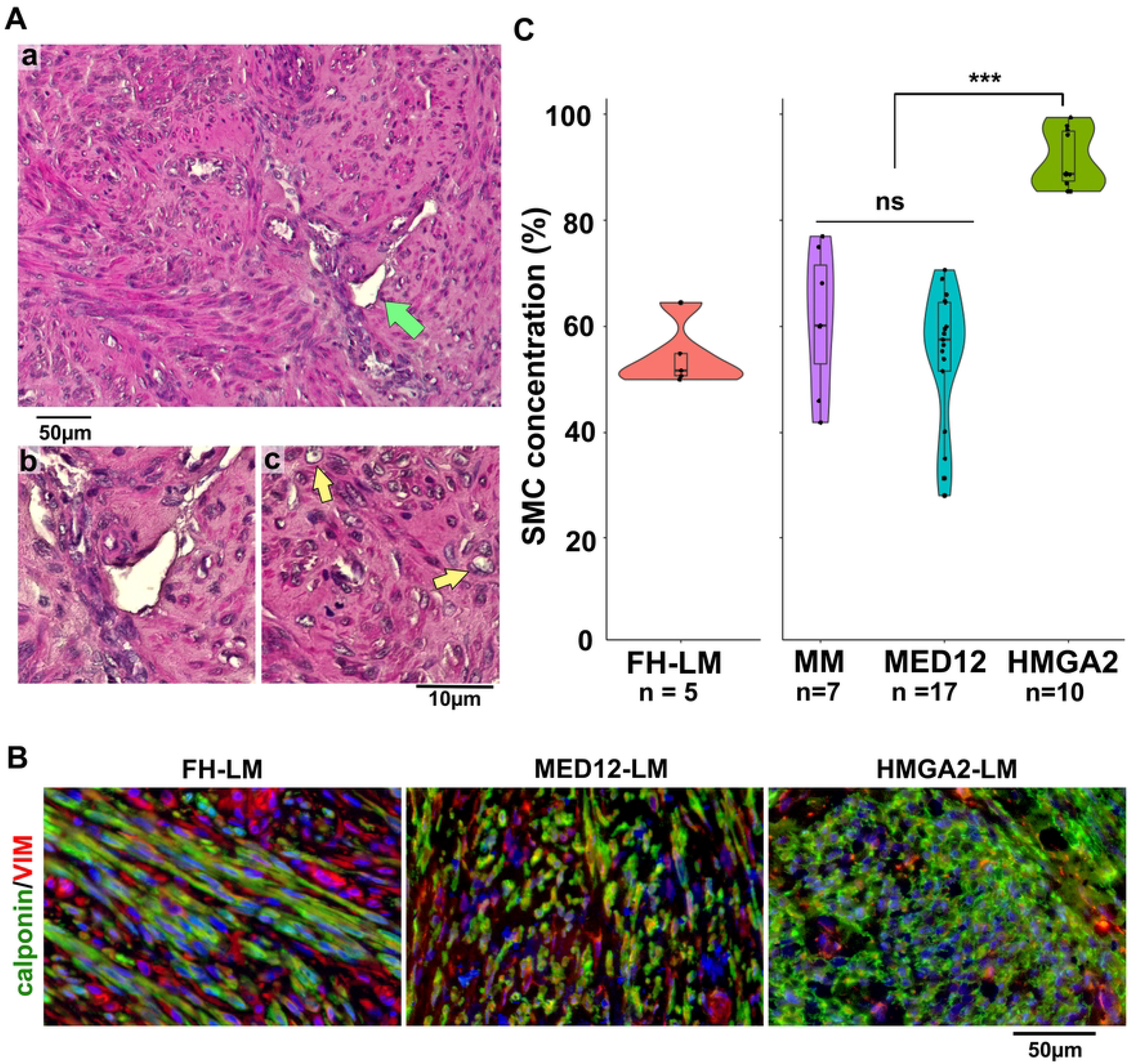
Histological characteristics of FH-LM A. The H&E staining detected morphological features characteristic for FH-LM in the FH-negative LM case, a staghorn-like vessel (arrow) (a) and enlarged nuclei with pseudoinclusions (c, arrows). B. IF detection of calponin (green) and VIM (red) in 3 LM subtypes. FH-LM and MED12-LM contained a substantial concentration of calponin-negative/VIM-positive cells (TAFs). C. Comparison of SMTC concentration among FH-LM, MED12-LM HMGA2-LM, and MM. Values of individual samples were plotted over a violin plot with an included boxplot. The original data for MM, MED12-LM, and HMGA2-LM (the right panel) were previously used in our study [7]. Each sample in these groups was derived from different patients. The five samples of FH-LM (the left panel) were derived from different parts of a single tumor. Statistical significance by ANOVA was indicated as *** p≤ 0.001 and ns (not significant)(p >0.05).

### FH expression patterns in MM and LM subtypes

In MM and FH-positive LMs, FH was predominantly expressed in SMCs, and FH signals in TAFs were visible only with an extended exposure time that saturated the signals in SMCs (Fig 3A). Co-localization with TOMM20, a mitochondrial marker, indicated that the high FH expression in SMCs reflected the high mitochondrial concentration (Fig 3B). FH was very low to undetectable in the FH-LM under the conditions used to detect FH in MM and other LM subtypes. Nonetheless, FH was detected in TAFs of the FH-LM when IF signals were detected with the extended exposure time (Fig 3A, arrow), suggesting that the loss of FH, the putative driver mutation, occurs only in SMTCs.

**Figure. 3.**
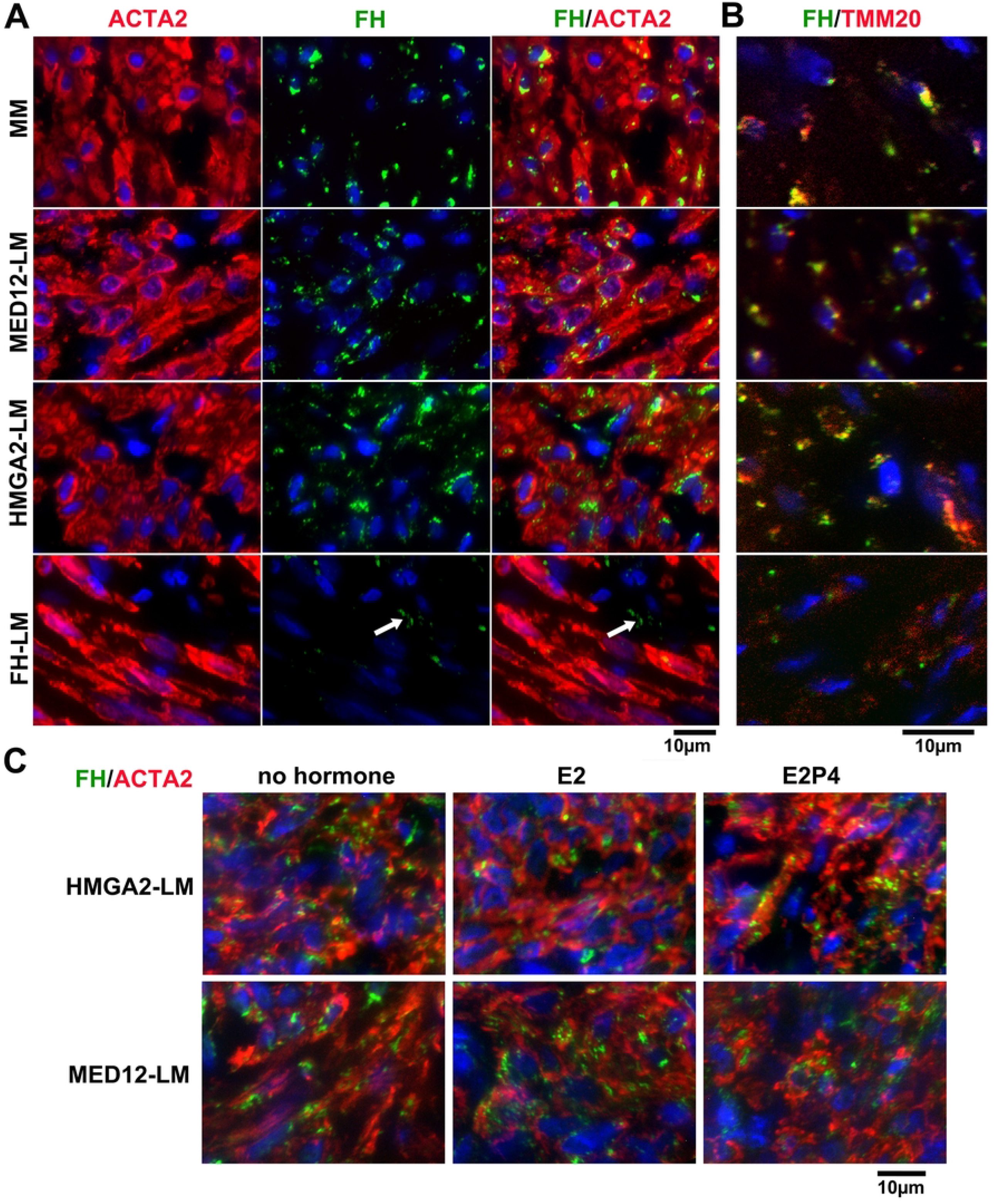
IF analysis of FH expression patterns in MM, MED12-LM, HMGA2-LM, and FH-LM subtypes. MM and LM samples were stained for FH (green) and either ACTA2 (red) (A and C) or TOMM20 (red) (B). A: FH was enriched in ACTA2-positive SMCs. B: Colocalization of FH and TOMM20 (yellow signal) indicated FH was expressed in mitochondria. C: FH expression in MED12-LM and HMGA2-LM PDXs treated with no hormone, E2, or E2P4 for 2 weeks [7]. In both LM subtypes, FH was constitutively expressed in SMTCs, and E2 and P4 had no detectable effects on the FH expression.

Since their growth depends on E2 and P4, the metabolism of MED12-LM and HMGA2-LM is likely regulated by ovarian steroids. Thus, we assessed the regulation of FH, a crucial enzyme in the TCA cycle, by E2 and P4 in LM PDXs. However, FH was constitutively expressed in SMTCs of MED12-LM and HMGA2-LM irrespectively of hormone treatments (Fig 3C).

### FH-LM overexpresses AKR1B10

Aldo-Keto Reductase Family 1 Member B10 (AKR1B10), an NADPH-dependent reductase that catalyzes the reduction of a wide variety of carbonyl-containing compounds, is often overexpressed in FH deficient tumors, including LM [4]. A recent study showed that AKR1B10 can be a specific and sensitive marker of FH-LM [24]. Our IHC assay confirmed the previous report and detected a high expression of AKR1B10 only in FH-LM but not in other LM subtypes (Fig 4A). Although an HMGA2-LM showed a weak signal (Fig 4A), it was not comparable to the FH-LM. In the FH-LM, AKR1B10 was overexpressed only on SMTCs (Fig 4B), suggesting that FH-deficiency upregulates AKR1B10 cell-autonomously.

**Figure. 4.**
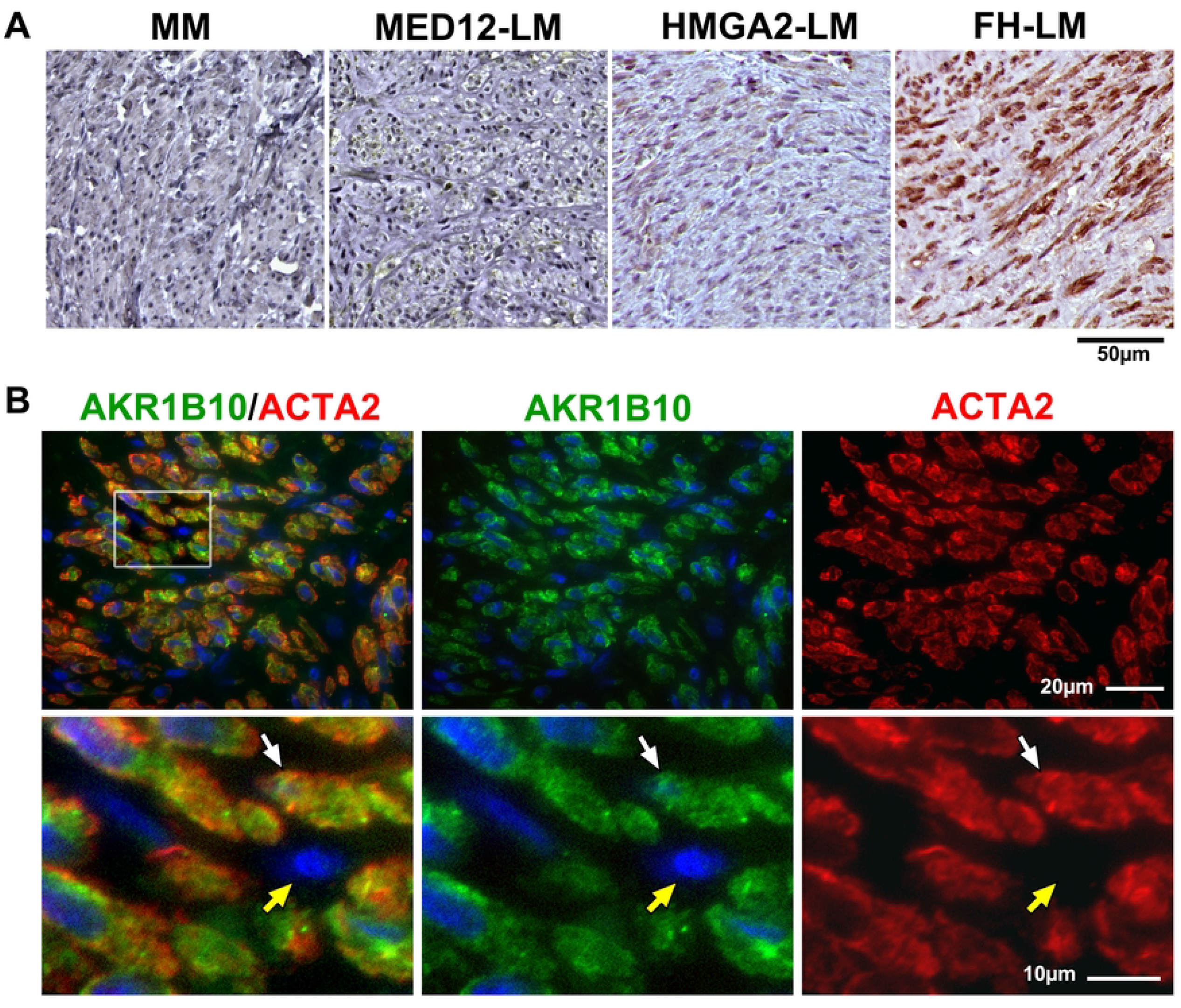
AKR1B10 expression patterns in LM subtypes A: Representative images of AKR1B10 IHC in MM, MED12-LM, HMGA2-LM, and FH-LM. AKR1B10 (brown) was highly expressed in the FH-LM, but not MM and other LM subtypes. B: Expression pattern of AKR1B10 (green) and ACTA2 (red) in FH-LM. AKR1B10 was overexpressed in SMTCs (white arrow) but not in TAFs (yellow arrow).

### E2 and P4 induce hypertrophy of SMTCs in FH-LM

FH-LM PDXs were subjected to hormone treatments, as shown in Figure 5A. Four weeks after grafting, a host mouse was euthanized to collect PDXs, and E2P4 pellets were replaced in other hosts. Two weeks after hormone pellet replacement, only the E2P4 group maintained tumor volume, and the PDXs of no hormone (NH) and E2 groups were significantly reduced in volume (Fig 5B, C). The regression of PDXs in NH and E2 groups was due to reduced SMTC size (Fig 6D). The increased cell density in NH and E2 groups (Fig. 5E) also supported that PDXs became smaller through cell size reduction but not cell death. These results indicate that FH-LM grows in response to E2 and P4 via SMTC hypertrophy, similarly to MED12-LM and HMGA2-LM.

**Figure. 5.**
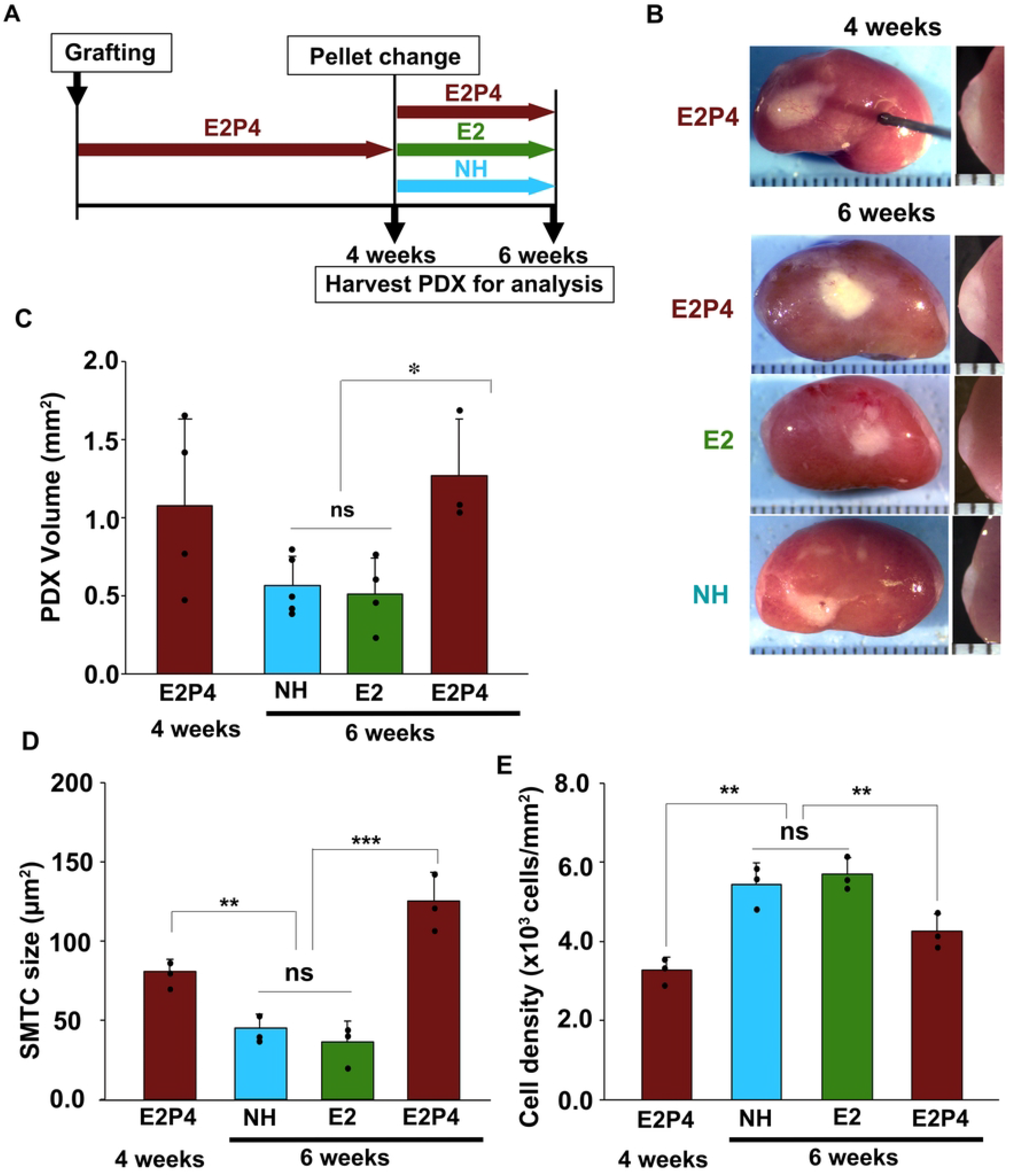
Hormonal response of FH-LM PDX A: Treatment schedule. PDXs were grown for four weeks in hosts supplemented with E2P4 and then subjected to one of three different treatments, no hormone (NH), E2 alone (E2), or E2P4 treatment, for two weeks (6 weeks after grafting). The PDXs were harvested for analyses at 4 and 6 weeks. B: FH-LM PDXs on the host kidney: whole grafts (left panel), grafts with kidney tissue were bisected to measure the height (right panel) [19]. C, D, and E: PDX volume (C), SMTC size (D), and cell density (E). At 6 weeks, PDX volume and SMTC size were significantly higher in the E2P4 group than in NH and E2P4 groups. In contrast, the cell density had a reverse correlation with PDX volume and SMTC size, indicating that the SMTC size is the primary factor that determines the tumor volume. Statistical significance by ANOVA was indicated as *P≤0.05, **P≤0.01, ***P≤0.001, and ns (P>0.05)

**Figure. 6.**
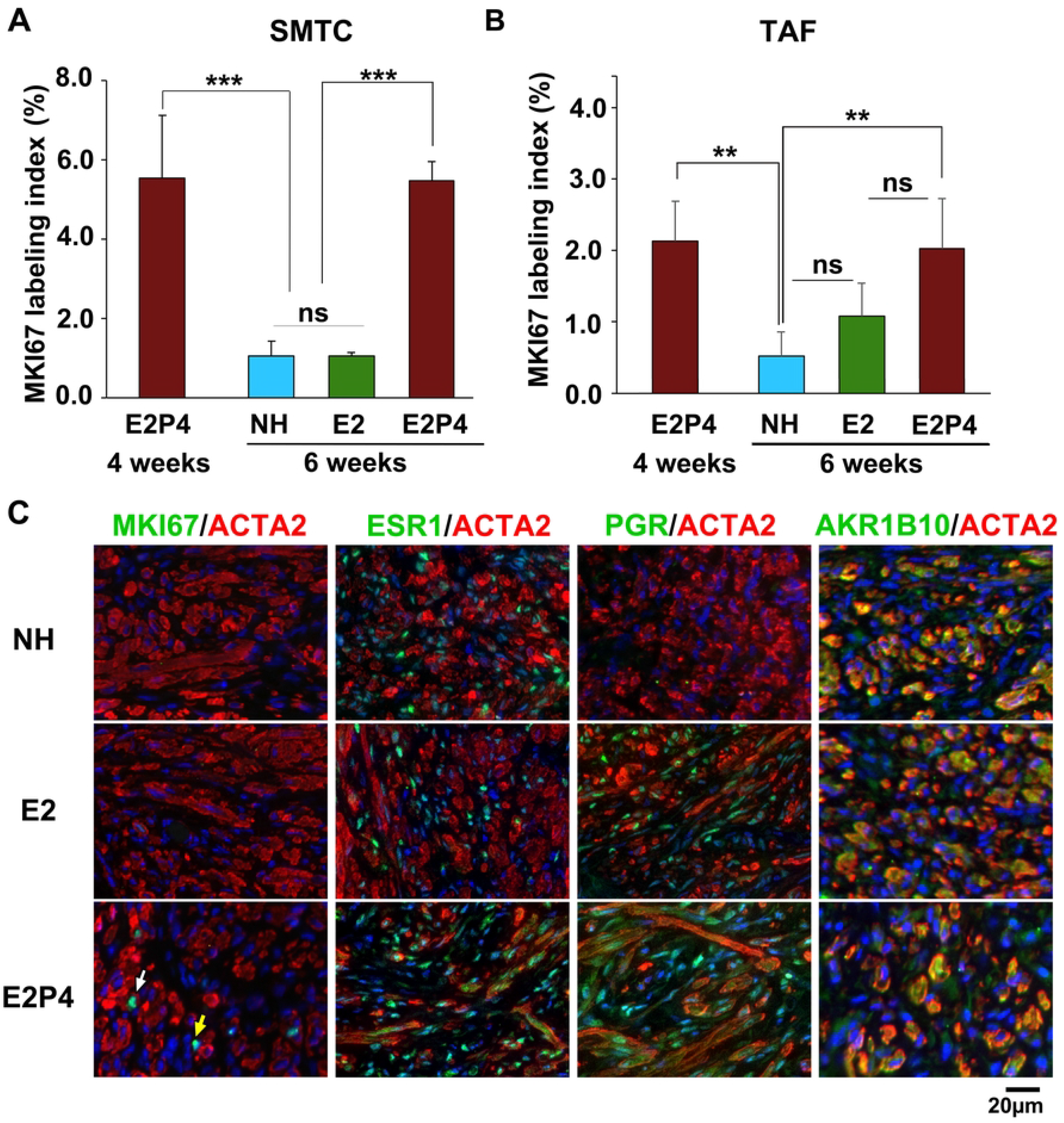
Hormonal regulation of cell proliferation and gene expression in FH-LM PDXs. A and B: MKI67 labeling index of SMTCs (A) and TAFs (B) in FH-LM PDXs. Statistical significance by ANOVA was indicated as *P≤0.05, **P≤0.01, ***P≤0.001, and ns (P>0.05). C: IF assay for ACTA2 (red) and MKI67, ESR1, PGR, or AKR1B10 (green).

### E2 and P4 stimulate cell proliferation in FH-LM

The proliferation activity in FH-LM PDXs was assessed by MKI67 IF. Like MED12-LM and HMGA2-LM, the MKI67 labeling index of FH deficient SMTCs was significantly higher in the E2P4 group than in NH and E2 groups (Fig 6A). Similarly, the MKI67 labeling index of TAF was significantly higher in the E2P4 group than in the NH group, indicating that the growth of TAFs in FH-LM depends on E2 and P4. Meanwhile, there was no significant difference in the proliferation rate of TAFs between E2 and E2+P4 groups or E2 and NH groups (Fig 6B), suggesting that in FH-LM, the growth of TAFs was most efficiently stimulated by the combination of E2 and P4, but E2 alone also has a weak growth-promoting effect.

### Hormonal regulation of gene expression in FH-LM

IF assays revealed that estrogen receptor α (ESR1) was expressed in both SMTCs and TAFs irrespective of hormone treatments. In contrast, the expression of progesterone receptor (PGR) depended on E2 in both SMTCs and TAFs, like MED12-LM and HMGA2-LM (Fig 6C). This result can explain why both E2 and P4 are required to stimulate the growth of FH-LMs.

Finally, we examined expression patterns of AKR1B10 in FH-LM PDXs. AKR1B10 was detected in SMTCs of all hormone treatment groups. Since AKR1B10 is constitutively expressed in FH-LMs, it can be an ideal surrogate marker for FH-deficiency.

## Discussion

Through a series of PDX studies, we have elucidated cellular mechanisms of LM growth and have further specified these mechanisms to LM subtypes. In this study, we characterized the response of FH-LM to ovarian steroids. Interestingly, the three most frequent LM subtypes share many biological characteristics despite the distinct gene expression profiles and unique morphological features: the driver mutations are exclusively present in SMTCs; the tumor volume increases by cell number (proliferation) and size (hypertrophy), and the growth of SMTCs depends on P4, but E2 is also required for PGR expression. Since MED12-LM, HMGA2-LM, and FH-LM together account for ∼90% of all LM cases, the current LM treatments targeting the hypothalamus-pituitary-ovary axis are effective for most LM patients.

FH deficiency is a putative driver of multiple human neoplasms, including highly aggressive renal cell carcinoma. Therefore, research on the pathogenesis of FH-deficient tumors is clinically significant even though they are rare. Unfortunately, the low incidence makes the research on FH-deficient tumors challenging [9-11, 25], and how FH deficiency promotes tumorigenesis remains elusive. It is especially intriguing why germline inactivation of an FH allele predisposes the carrier to certain types of neoplasms, even though FH is ubiquitously expressed as a vital enzyme in the mitochondrial respiratory chain [26].

FH deficiency results in reduced 2-oxoglutarate (2-OG)-dependent dioxygenase (2-OGDD) activity. The 2-ODGGs are a large group of enzymes that catalyze hydroxylation reactions on various substrates (e.g., protein, nucleic acid, lipid, and metabolic intermediate), producing CO2 and succinate. The activity of 2-OGDD depends on the intracellular ratio of 2-OG to inhibitors such as fumarate, succinate, and 2-hydroxyglutarate. It has been proposed that hypoxic responses triggered by reduced 2-OGDD activities contribute to the pathogenesis of FH-deficient tumors [27, 28]. In addition, the accumulation of fumarate inhibits 2-OG-dependent histone and DNA demethylases [27-29]. Thus, it has also been implied that FH deficiency promotes neoplastic transformation by epigenetic reprogramming. Furthermore, the accumulation of S-(2-succino)-cysteine (2SC) covalent modifications [25] may also play a role in the formation of FH-deficient tumors as epigenetic modifiers. The mechanisms mentioned above have been examined primarily in renal cell carcinoma. However, while renal cell carcinoma grows cell-autonomously, FH-LMs depend on E2 and P4 in their growth. Thus, whether FH-deficient LMs and renal cancers share molecular pathogenesis is unclear.

Accordingly, the mechanisms through which FH deficiency causes tumors should be studied in the myometrium. However, the primary cell culture is unsuitable for studying the pathogenesis of FH-LM that contain a high concentration of TAFs. Thus, PDX studies with additional FH-LM cases are essential.

Finally, our PDX studies determined the P4-dependency of the three most prevalent LM subtypes in their growth [5-8, 19]. However, ∼10 % of LMs lack hallmark mutations of MED12-LM, HMGA2-LM, and FH-LM and show unique molecular signatures [4]. The growth control of these rare LMs may be distinct from the three major LM subtypes. In this respect, epidemiological studies showed the essential role of ovarian steroid in the pathogenesis of LM, but the effect of E2 and P4 could not be assessed separately. Accordingly, some rare LM subtypes may be stimulated by E2 alone and thus unresponsive to SPRM treatments. Thus, understanding the growth characteristics of rare LM subtypes, including COL4A5/6-LM, is of utmost significance to designing treatment strategies for LMs.

## Acknowledgements

We thank The Genomics Shared Resource (GSR) at The Ohio State University Comprehensive Cancer Center, and The Tissue Procurement Service Laboratory at The Ohio State University Wexner Medical Center.

## Grant Support

Research reported in this publication was supported by the National Institutes of Health [R21HD102897 to T.K.], The Ohio State University Comprehensive Cancer Center and the National Institutes of Health [P30 CA016058].

## Notes

### Competing Interest Statement

The authors have declared no competing interest.

